# The use of principal component and factor analysis to measure fundamental cognitive processes in neuropsychological data

**DOI:** 10.1101/2021.11.10.468133

**Authors:** Christoph Sperber

**Affiliations:** University of Tübingen

**Keywords:** Principal component analysis, PCA, factor rotation, dissociation, de-noising, cognitive assessment, association problem, deficit, lesion, stroke

## Abstract

For years, dissociation studies on neurological single cases with brain lesions were the dominant method to infer fundamental cognitive functions in neuropsychology. In contrast, the association between deficits was considered to be of less epistemological value and even misleading. Still, principal component analysis (PCA) – an associational method for dimensionality reduction – recently became popular for the identification of fundamental functions. The current study evaluated the ability of PCA and factor analysis (FA) to overcome the association problem in behavioural data of neurological patients and to identify the fundamental variables underlying a battery of measures. Synthetic data were simulated to resemble neuropsychological data with typical dissociation patterns and, thereby, typical patterns of dependence between variables. In most experiments, PCA and FA succeeded to measure the underlying target variables with high up to almost perfect precision. However, this success was fragile and relied on the success of factor rotation, which failed its intended purpose when no test scores existed that primarily measured each underlying target variable. Further, commonly used strategies to estimate the number of meaningful factors appear to underfactor neuropsychological data, thereby consistently underestimating the dimensionality of the data. Finally, simulations suggested a high potential of PCA to denoise data, with factor rotation providing an additional filter function. This can be invaluable in neuropsychology, where measures are often inherently noisy, and PCA can be superior to common compound measures, such as the arithmetic mean, in the measurement of variables with high reliability. In summary, PCA and FA appear to be powerful tools in neuropsychology that are well capable to infer fundamental cognitive functions with high precision, but the typical structure of neuropsychological data and limited informative value of associations in neuropsychology place clear limitations and a risk of a complete failure on the methods.

## 1 Introduction

The study of neurological single cases has been a powerful catalyst for cognitive research. Landmark cases like the aphasia patient reported by Paul Broca (see Dronkers et al., 2007) created the early foundation of cognitive science. Later, after a period with a focus on group studies, the single case method re-emerged in the 1960s to be considered by some as “the most powerful empirical procedure for making inferences to normal function” (p. 15 in Shallice, 1988). Especially the dissociation of cognitive functions was considered pivotal to the scientific value of single cases, and the dissociation method was widely used to identify fundamental mental processes underlying behaviour. This was mirrored by an extensive theoretical background on the dissociation method. Different subtypes of dissociations were defined (Shallice, 1988; Crawford et al., 2003), sophisticated statistics to identify dissociation types were developed (e.g. Crawford & Garthwaite, 2002), and an entire special issue in the journal Cortex was dedicated to the critical discussion of the method (Dunn & Kirsner, 2003). Nevertheless, the popularity of single case studies drastically declined in the last two decades and most of cognitive neuroscience shifted towards group studies.

Neuropsychological group studies require different analysis strategies. Among them, principal component analysis (PCA) became popular, which is an adaptive exploratory method that does not make any distributional assumptions (Jolliffe & Cadima, 2016). Principal component analysis rotates the axes of a data set in a way that the first few axes explain as much of the given variance as possible. For example, imagine the scores in six language tests that were measured in a sample of stroke patients. A PCA might rotate the six axes (one for each original language score) so that only two variables describe most of the variance in the data, and one might interpret these two variables to represent fundamental language functions. Likewise, but to a much lesser extent, factor analysis (FA) has been used for the same purpose. Factor analysis estimates an ideally low-dimensional factor model to explain the data. Although FA is therefore mathematically different from PCA, conceptually it is used in a very similar way.

The identification of fundamental cognitive processes is a common rationale behind the use of PCA on neuropsychological data from patients with brain damage (e.g., Verdon et al., 2010; Randerath et al., 2011; Butler et al., 2014; Chechlacz et al., 2014; Mirman et al., 2015; Chen et al., 2016; Fridriksson et al., 2016; Halai et al., 2017; Aguilar et al., 2018; Alyahya et al., 2018; Tochadse et al., 2018; Schmidt et al., 2022). The same approach has also been used more descriptively to characterise the complexity of neuropsychological test profiles (e.g., Azouvi et al., 2002; Zandieh et al., 2012; Corbetta et al., 2015; Timpert et al., 2015; Sperber & Karnath, 2016; Bisogno et al., 2021). However, there is often no clear border between the use of principal component analysis to identify fundamental cognitive functions from test profiles or to simply describe test profiles. Some studies used principal component analysis to investigate extensive test batteries that covered a large number of functional domains (Corbetta et al., 2015; Bisogno et al., 2021). They found that neuropsychological pathology can largely be explained with only three factors and, besides this merely descriptive finding, they concluded that the factors correspond to the connectomic organisation of the brain. What is particularly enigmatic about these findings is that the methods behind them do not differ substantially from the studies mentioned above. The same method is interchangeably used to either infer fundamental cognitive functions from a test battery across a single modality (such as language deficits) or clusters of deficits from a behaviourally multimodal test battery.

Finally, some studies used PCA to obtain one single score from a set of highly similar measures and used the first component as the study’s main measure (Rondina et al., 2016; Rondina et al., 2017). A reason behind this can be the intention to create a composite score from differently scaled measures. The computation of a mean score would be invalid in such a situation, but a PCA could still succeed to create a valid composite measure. Second, and although not explicitly stated in these publications, PCA might be used to de-noise data. If each variable measures the same fundamental quantity with noise, the first factor in the PCA should capture this variable, while any succeeding factors might capture a small portion of noise. Hence, the first factor might provide a de-noised measure of the fundamental quantity. This rationale underlies the common use of PCA in image denoising (Zhang et al., 2010).

All these strategies behind componential analysis in neuropsychological data sound highly appealing for the study of cognition and its pathology. However, few exceptions aside (e.g. Halai et al. 2017), many studies only shallowly described the theoretical background behind their method. Extensive theoretical background, critical discussions and evaluations as available for the dissociation method are lacking. Further, the exact methodology differs between studies in the choice of factor rotation strategies (compare e.g. Halai et al. 2017 and Bisogno et al. 2021), and a systematic comparison or a theoretical foundation that compares orthogonal versus oblique factor rotation does not exist. The lack of a deeper theoretical background might be especially problematic as PCA accounts for the association between variables. It is long known that the unidimensionality of a set of behavioural measures does not imply that only a single process underlies them, as the processes could function in unison (Bejar, 1983). An association problem in neuropsychology adds another layer to this issue, as processes might be damaged in unison, independent of their functional relationship. A prime example of this association problem is the Gerstmann syndrome (see Shallice, 1988), in which four apparently unrelated deficits were erroneously combined into one syndrome because of their consistent common occurrence. It was not until much later that the view gained acceptance that these are cognitively non-overlapping deficits whose neuronal correlates lie in close proximity and are usually damaged together by typical stroke. Due to this association problem, neuropsychology focussed on the dissociation method for such a long time. Therefore, when deficits in cognitive functions occur highly consistently, associational computational methods such as PCA are unable to differentiate them. However, even for situations where co-occurrence is not perfectly consistent, it was found that lesion mapping of principal components obtained from neuropsychological data can result in artefacts (Sperber et al., 2020). In summary, it remains an open question how well PCA and FA perform in the face of the association problem in neuropsychology, and if and how well they can infer fundamental cognitive functions from associations in group data.

In the present study, I evaluated the adequacy and efficiency of principal component and factor analysis in the identification of fundamental cognitive functions underlying multiple neuropsychological measures. Through simulations of synthetic data, perfectly transparent conditions resembling neuropsychological data were generated, including typical dissociation patterns between deficits, i.e. typical dependencies between variables. These allowed a systematic evaluation of these methods in neuropsychology: what does principal component or factor analysis do if two distinct, hidden cognitive processes are measured with a test battery? What if these processes are not independent, but they systematically co-occur with occasional dissociations, as typical for neuropsychological data? What does it do if a variable is assessed with multiple highly noisy items?

## 2 Methods

### 2.1 General methods

The study is entirely based on synthetic data that I simulated to resemble idealised neuropsychological data. Only the use of synthetic data allowed to test componential analyses in a setting where hidden fundamental cognitive processes and their relation to each other were perfectly known. Analyses were scripted with MATLAB 2021a. All scripts are available in the online materials.

The general study design is illustrated in Figure 1. Possible specifications and modifications of this design are described in the text section of each experiment. Each experiment consisted of 100 simulations, and each simulation was run for 100 patients, which is a large, but realistic sample size for neuropsychological data. In each experiment, one or two target variables were simulated. These were thought to represent fundamental cognitive functions that a neuroscientist wants to infer from a battery of test scores as precisely as possible by the use of principal component analysis. Within each sample of 100 patients and for each target variable, 50 patients were randomly chosen to have a deficit and 50 to not have a deficit. For patients with the deficit, the target variable was randomly picked from a normal distribution with μ = 1 and σ = 0.4, and for patients without the deficit with μ = 0 and σ = 0.4. Any values further away from the mean than 0.5 (i.e. 1.25*σ) were re-rolled, hence the value 0.5 was a clear cutoff between pathological and normal scores. This reduced the actual standard deviation of each group from 0.4 to ∼0.26. Based on the target variables, a test battery of six scores was simulated. Each test score was generated additively from uniformly distributed noise and a contribution of the target variables. I set the noise function to pick values between 0 and 0.4. With the inclusion of noise, I intended to create synthetic data that resemble actual neuropsychological data, which are usually noisy. This procedure was previously explained and empirically derived (Pustina et al., 2018).

**Figure 1.**
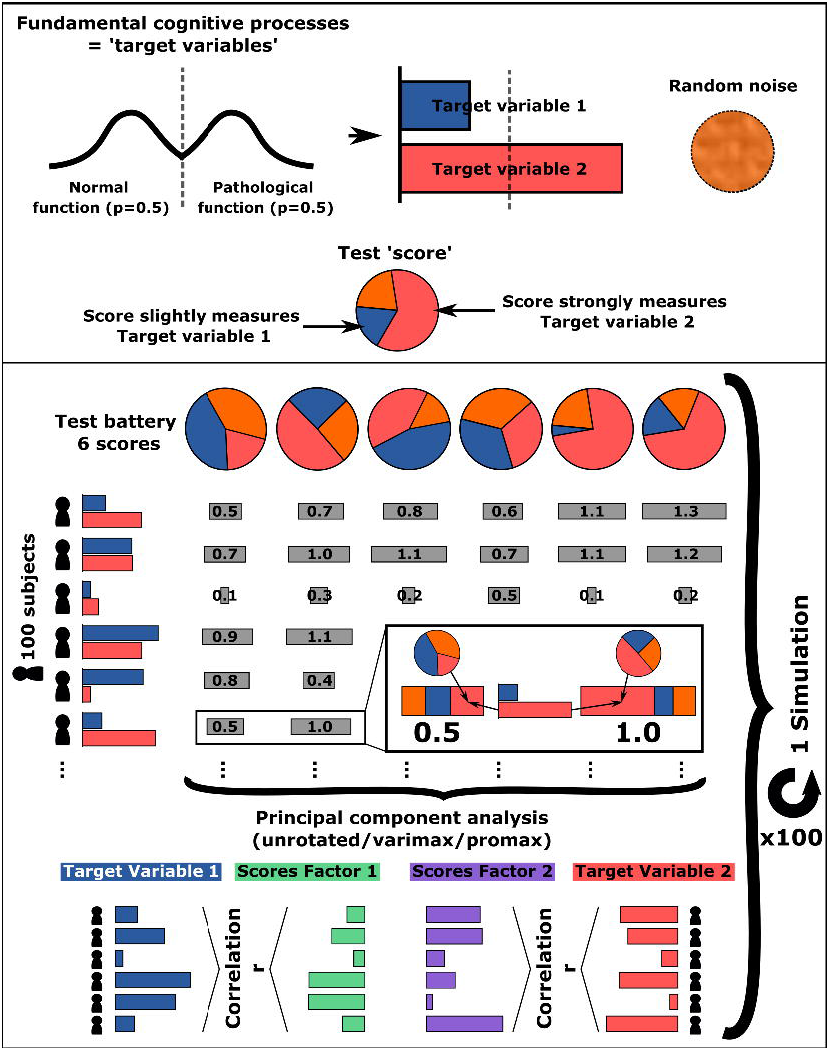
Study design - Experiment 1a. Illustration of the study design in experiment 1a. The design of all other experiments resembled this design, and the relevant changes are described in each experiment’s method section.

Principal component analysis was run on the test battery of six scores for each simulation in Matlab with the ‘pca’ function, which centres the data and uses singular value decomposition. Factors were rotated by the ‘rotatefactors’ function for orthogonal varimax or oblique promax rotation using default options, except for an increased iteration limit. I always rotated the first two factors, independent of any criteria to define an optimal number of factors. The main dependent variables were i) the Pearson correlation r of the hidden target variables with the first two factors found by the PCA and ii) the variance explained by these factors. If only one target variable existed, the first dependent variable was its correlation with the first factor. When two target variables existed, it was not known in advance which of the two factors better assesses each target variable. Therefore, I always chose the factor-variable pair with the higher correlation. However, this procedure would miss cases in which the PCA failed to align the first two factors with the target variables as intended. In such a case, either one factor would correlate higher with both target variables, or both factors would correlate higher with the same target variable. This would indicate failure of the methodology, hence these cases were identified and excluded from further analyses. If present, the number of such cases is reported. The remaining results for the unrotated PCA versus orthogonal varimax rotated PCA versus oblique promax rotated PCA were compared by Monte Carlo permutation-based two-sample t-tests with 100.000 permutations that were corrected for multiple comparisons by the Bonferroni algorithm. As the groups were not fully paired I performed independent tests. The detailed descriptive and statistical results are reported in tables 1-3, and within the text I only report abbreviated results to increase readability. The explained variance was qualitatively evaluated by reference to the Kaiser criterion (Kaiser, 1960). The Kaiser criterion provides a strategy to estimate the number of factors to be retained and rotated in a PCA, and it is popular in neuropsychology (e.g. Corbetta et al., 2015; Halai et al., 2017). Following this criterion, only factors with an eigenvalue >1 are retained, i.e. factors that explain more variance than a single original variable. With 6 variables in each simulation run, this corresponds to a factor that explains more than 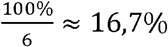 of the variance. Hence, a factor that lies below this value would not be identified as meaningful. If only one factor surpasses this value in the presence of two hidden target variables, this would again indicate a failure of the methodology. However, several other strategies to estimate the number of factors exist. Therefore, and to make different analyses comparable, I retained these cases in the statistical analysis.

**Table 1.** Detailed results of experiments 1-3 with PCA. The first column reports the variance explained by the first (and second) factor of the PCA. The following three columns report the correlation of the first (and second) factor with the corresponding target variable. Simulations in which the first two factors failed to align with the target variables are reported in the text and were not considered here. The last three columns show the results of the statistical comparison between the three different PCA solutions. Values are mean±standard deviation. Significant results are highlighted in bold. UR = unrotated; VM = varimax; PM = promax.

| Experiment | Expl. | Correlation r |  |  | Significance p |  |  |
| --- | --- | --- | --- | --- | --- | --- | --- |
|  | Variance in %<br>1<sup>st</sup>/(2<sup>nd</sup>)<br>Factor | Factor(s)-Target Variable(s) |  |  | (Bonferroni-corrected) |  |  |
|  |  | Unrotated | Varimax | Promax | UR-VM | UR-PM | VM-PM |
| 1a | 77.6±6.8 /<br>18.5±7.0 | 0.84±0.08 | 0.93±0.06 | 0.95±0.05 | < .**.001** | < .**.001** | < .**.001** |
| 1b | 61.4±9.5 /<br>30.5±9.6 | 0.93±0.07 | 0.97±0.03 | 0.97±0.03 | < .**.001** | < .**.001** | = 1.00 |
| 2a | 85.0±4.8 /<br>11.5±4.8 | 0.83±0.10 | 0.95±0.04 | 0.96±0.04 | < .**.001** | < .**.001** | < .**.05** |
| 2b | 65.4±7.5 /<br>26.3±7.6 | 0.87±0.09 | 0.97±0.03 | 0.97±0.03 | < .**.001** | < .**.001** | = 1.00 |
| 3 low Noise | 68.8±4.6 | 0.96±0.01 | 0.93±0.03 | 0.92±0.05 | < .**.001** | < .**.001** | < .**.01** |
| 3 med Noise | 52.9±9.3 | 0.91±0.04 | 0.93±0.02 | 0.93±0.03 | < .**.001** | < .**.01** | = .56 |
| 3 high Noise | 33.7±4.3 | 0.76±0.06 | 0.92±0.04 | 0.90±0.05 | < .**.001** | < .**.001** | = .16 |

**Table 2.**
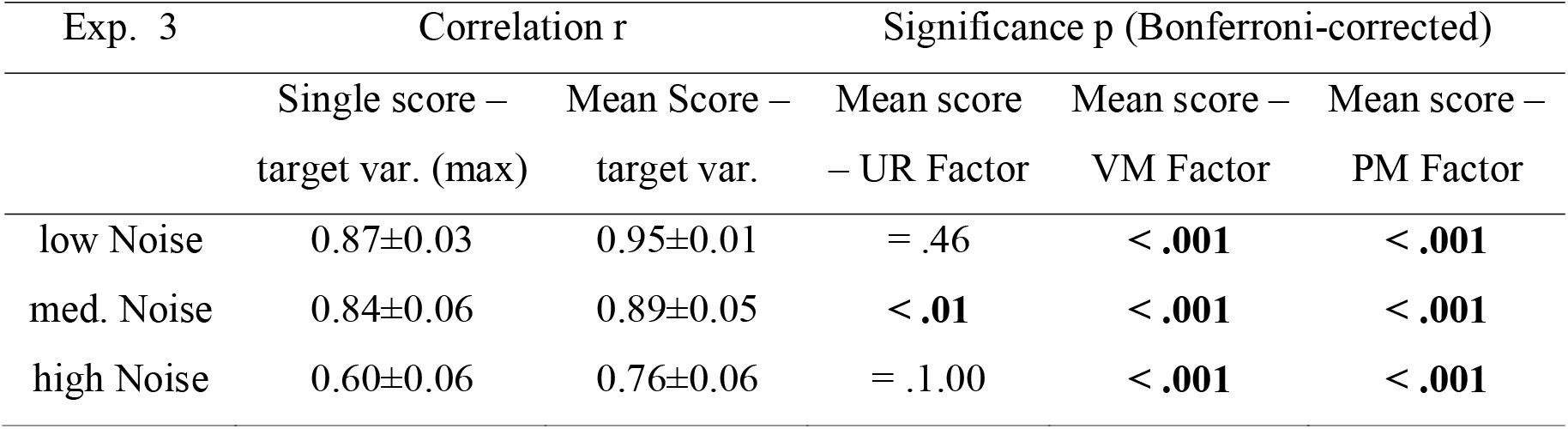
Additional results of experiment 3. The first two columns show the correlation between the target variable and a) the single score out of all six with the highest correlation and b) the mean score of all six scores. The remaining columns show the statistical difference between the latter and the correlation of the target variable with the first PCA factor. In other words, the statistical results indicate if the ability of a PCA solution to measure the target quantity was different from the mean score’s ability to do so. The corresponding correlations between the target variable and the PCA factors for the unrotated (UR) solution and the rotated varimax (VM) and promax (PM) solutions are shown in Table 1. Values are mean±standard deviation. Significant results are highlighted in bold.

**Table 3.** Detailed results of the additional posthoc experiments with PCA. Detailed results for experiments 2c and 2d. See the legend of table 1 for more information. Values are mean±standard deviation. Significant results are highlighted in bold.

| Experiment | Expl. | Correlation r |  |  | Significance p |  |  |
| --- | --- | --- | --- | --- | --- | --- | --- |
|  | Variance in %<br>1<sup>st</sup>/(2<sup>nd</sup>)<br>Factor | Factor(s)-Target Variable(s) |  |  | (Bonferroni-corrected) |  |  |
|  |  | Unrotated | Varimax | Promax | UR-VM | UR-PM | VM-PM |
| 2c | 75.6±4.8 /<br>16.0±4.0 | 0.83±0.18 | 0.97±0.03 | 0.97±0.03 | < **.001** | < **.001** | = 1.00 |
| 2d | 81.4±4.7 /<br>9.0±2.1 | 0.84±0.13 | 0.88±0.10 | 0.88±0.10 | < **.05** | < **.05** | = 1.00 |

Factor analysis was computed using the ‘factoran’ Matlab function with maximum likelihood estimation. As with principal component analysis (PCA), factors were rotated either with orthogonal varimax or oblique promax rotation. With a predefined random number generator seed, I ensured that the factor analyses were applied to the same simulated behavioural data as principal component analysis. Again, two factors were rotated in all analyses, independent of any criteria to select the number of factors. Additionally, the Kaiser-Meyer-Olkin Index (Kaiser & Rice, 1974) was computed. This measure ranges between 0 and 1 and indicates if a dataset is suitable for factor analysis. Commonly, values above 0.5 are expected to be required for adequate factor analysis. Compared to PCA, the use of factor analysis on behavioural data in neurological patients is rare. For this reason, and to increase the article’s readability, the main manuscript only reports PCA results, and the results of the factor analyses are reported in the supplementary.

### 2.2 Simulation experiment 1a – two independent cognitive functions

A popular rationale for the application of PCA on neuropsychological data is the identification of fundamental cognitive functions. Imagine, for example, that we measure attention with multiple tests. Each test score assesses spatial and non-spatial attention to different degrees. With the application of PCA, we could aim to directly and exclusively assess the deficit in spatial attention respectively the deficit in non-spatial attention.

In this first experiment, the deficits represented by the target variables were independently chosen. The average correlation between the two target variables across all simulations was r = 0.01 (SD = 0.09). I simulated each score in the test battery based on a random ratio of the first and the second target variable. This ratio was set constant for each score across all patients within a simulation. In the above example, this would correspond to test items that are, to a certain degree, consistently affected both by spatial attention and non-spatial attention.

#### Results

The first factor explained on average 77.6%, and the second factor 18.5% of the variance. In 42 out of 100 simulations, the second factor did not exceed the Kaiser criterion, and thereby, if strictly following this criterion, the PCA failed to find a two-factorial solution. In twelve simulations with unrotated factors and seven simulations each after varimax and promax rotation, the first two factors were not aligned with one target variable each. Nonetheless, after the removal of these cases, the PCA factors correlated highly with the target variables. The unrotated solution (r = .84) was worse than both rotated solutions, and promax rotation (r = .95) was slightly superior to varimax rotation (r = .93). Detailed results are reported in table 1.

Simulations in which rotated factors failed to describe one target variable each consisted only of cases in which both factors correlated higher with the same target variable. For example in the 86^th^ simulation and after varimax rotation, the first PCA factor correlated higher with the second variable (r = .98) than the first (r = .28), and likewise did the second PCA factor correlate higher with the second variable (r = .85) than the first (r = .59). The reversed type of factor-variable misalignment – a factor that correlates higher with both target variables – was only found for unrotated solutions. For example in the 13^th^ simulation, the first factor correlated higher with both target variables (r = .75; r = .72) than the second factor (r = .65; r = .66).

#### Discussion

In this first experiment, PCA showed variable performance. On the one hand, it failed in a few simulations to align the first two factors with the target variables, and in 42% of simulations, the Kaiser criterion underestimated the dimensionality of the target variables by suggesting a one-factorial solution. On the other hand, the remaining rotated solutions precisely captured the target variables as intended.

With this experiment, I intended to set an optimistic baseline for the capability of PCA to identify the target variables in the present simulation setting. The difficulties to identify a two-factorial solution were unexpected but explainable in hindsight. While both target variables are independent, the six test scores are not. All test scores measure both target variables to some degree, and hence they correlated by r = .73 (SD = .22). In the end, with the battery of six correlating scores, a single factor was often sufficient to explain a large share of the total variance, and the Kaiser criterion failed to identify the presence of a second underlying target variable. This unexpected finding points at a highly relevant issue: even if fundamental quantities are independent, measures assessing these quantities can be dependent. And when all measures are dependent, a single factor can explain their shared variance.

##### 2.2.1 Simulation experiment 1b – two independent cognitive functions and independent test scores

In this variation of experiment 1a, I ensured that not only the target variables but also the scores that were created from them were independent. The only change compared to experiment 1a was that, out of the six test scores in each simulation, three scores only assessed the first target variable, and the three other scores only the second. The correlation across all items was r = .27 (SD = .37), however, across the two groups of three items each, items did not correlate (r = .01; SD = .09).

#### Results

The first factors explained on average 61.4% and 30.5% of the variance. The second factor did not exceed the Kaiser criterion in 12 out of 100 simulations. This time, the first two PCA factors were always aligned with one target variable each. Promax and varimax rotation (both r = .97) performed equally and better than the unrotated solution. Detailed results are reported in Table 1

#### Discussion

With independent target variables and measures that each assessed only one target variable, PCA performed very favourably. The first two factors – both rotated and unrotated – always aligned with the target variables, and rotated solutions measured the target variables almost perfectly. Still, in a few cases, the Kaiser criterion failed to identify both factors.

### 2.3 Simulation experiment 2a – two dependent cognitive functions

The simulations in experiments 1a and 1b included a tacit assumption that makes the simulated fundamental cognitive deficits, i.e. the target variables, unrepresentative of most actual deficits: they were statistically independent. In this experiment, the only change over experiment 1a was that I did not simulate each target variable independently, but based on a pre-defined dissociation pattern. In experiment 1a, 50 patients were randomly chosen to suffer from a deficit. On average, this resulted in 25 patients with both deficits, 25 without the deficit and 25 each with only one of the deficits. In experiment 2a, I randomly picked 35 patients to suffer both deficits, 35 to suffer no deficit, and 15 patients to suffer only one deficit. This resulted in target variables that were moderately correlated by r = .32 (SD = 0.07). The six test scores correlated strongly with each other with r = .82 (SD = .14)

#### Results

The first factors explained on average 85.0% and 11.5% of the variance. The second factor did not exceed the Kaiser criterion in 85 out of 100 simulations, and the factors failed to describe one target variable each in 86 out of 100 simulations with the unrotated solution, in eleven simulations with varimax rotation and seven simulations with promax rotation. Again, rotated solutions outperformed unrotated ones with a small advantage of promax rotation (r = .96) over varimax rotation (r = .95). Detailed results are reported in table 1.

#### Discussion

The performance of PCA to identify statistically dependent fundamental cognitive functions seemed to be limited in this experiment. In a large majority of the simulations, the Kaiser criterion failed to identify the two-factorial data structure, and the unrotated solutions failed most often to align with the target variables. Nonetheless, most rotated solutions succeeded to capture the target variables, and the successful solutions precisely assessed the target variables.

##### 2.3.1 Simulation experiment 2b – two dependent cognitive functions and independent test scores

This experiment replicated experiment 1b with the same modification that was introduced in experiment 2a. Target variables followed a dissociation pattern (35-15-15-35) and, therefore, were statistically dependent.

#### Results

The first factors explained on average 65.4% and 26.3% of the variance. The second factor did not exceed the Kaiser criterion in 16 out of 100 simulations, but the first two factors always aligned with one target variable each in rotated solutions. The unrotated solution, however, failed in 25 out of 100 cases. Both rotated solutions outperformed the unrotated solution with an equally high correlation with the target variables (both r = .97). Detailed results are reported in table 1.

#### Discussion

Like in experiments 1a and 1b, PCA performed much more favourably when the scores measured only one target variable each, and thus not all scores were correlated. Rotated solutions performed as intended in all simulations and almost perfectly assessed the target variables.

### 2.4 Simulation experiment 3 – one cognitive function measured with high noise

One potential application of PCA on neuropsychological data is the de-noising of a single measure. Imagine that we measure primary motor deficits of the upper limb with multiple tests. No test provides perfect reliability and the psychological assessment of neurological patients is subject to many adversities. Hence, each single test score will only provide a noisy measure of primary motor deficits. We might apply PCA to measure primary motor deficits from several noisy scores with the rationale that the first PCA factor will mainly capture primary motor deficits, while all succeeding factors capture a small portion of noise. Importantly, the use of PCA would only be generally beneficial if it outperforms the mean score of all test scores.

For each simulation, I created a single target variable that was measured by all six test scores with noise. This procedure was repeated across three conditions with different noise levels. In detail, for each score, a noise proportion n was chosen. Each test score t was computed additively from a uniformly random value m between 0 and 1 and the target variable v by the formula t= n*m + (n-1)*v. The noise proportion n was chosen uniformly random. In the low noise condition, it was chosen between 0.5 and 0.7, in the medium noise condition between 0.5 and 0.9, and in the high noise condition between 0.7 and 0.9. I chose these values after piloting simulations to produce scores with varying degrees of medium to high correlations with the target variables. The average correlations of the scores with the target variables were r = .78, r = .62, and r = .43. In line with the other experiments, I also performed a factor rotation. I always rotated two factors, even though a uni-factorial solution might better describe the data. The reason is that a single factor simply cannot be rotated in isolation.

#### Results

The mean of all six test scores performed numerically better in capturing the target variable than the single best measure in all noise conditions. Hence, I chose the mean score as a reference to evaluate the performance of PCA. The first factor explained between 33.7% of the variance in the high noise condition and 68.8% in the low noise condition. The unrotated PCA solution performed similarly to the mean score; it was minimally better only in the medium noise condition. Across medium and high noise conditions, rotated factor solutions were significantly better than the mean of all six scores. Only in the low noise condition, the mean score was superior to rotated PCA solutions. However, this was not due to a failure of the PCA. The rotated scores strongly correlated with the target variable (r = .92/ r = .93). The mean score, however, was even better suited to explain the target variable in the low noise condition (r = .95), but its performance decreased with increasing noise to r = .76. Detailed results are reported in table 1, and additional results are reported in table 2.

To obtain a better understanding of this unexpected finding, I explored the PCA coefficients of the first factor in the high noise condition, where the superiority of rotated solutions was most striking. First, for each simulation, I identified the single score out of six that provided the worst measure of the target variable, i.e. which correlated the lowest with the target variable. The average coefficient for the unrotated solution was 0.24, for the varimax solution 0.05, and the promax solution 0.04. Next, I repeated the same procedure with the single score out of six that provided the best measure of the target variable. The average coefficient for the unrotated solution was 0.51, for the varimax solution 0.45, and the promax solution 0.44. In summary, the rotated factors appeared to depend less on the noisiest scores than unrotated solutions.

#### Discussion

Not surprisingly, the mean score as a compound measure of multiple noisy individual measures was better suited to assess a quantity than each measure in isolation. The first factor of unrotated PCA solutions was on a level with the mean score or, at best, marginally superior. This illustrates that PCA can be used to create a compound score, which could be a valuable perk whenever variable scaling impedes the computation of central tendency measures. However, when a mean score can be computed, an unrotated PCA might not provide any advantages over the mean score.

A highly interesting finding was that rotated PCA solutions were well capable to assess the target variable in all noise conditions, and even when individual measures were highly noisy. In the medium and high noise conditions, rotated factor solutions outperformed the mean score and the unrotated solution. Surprisingly, they captured the target variable with high precision. Factor rotation commonly aims to find a simple structure (Kaiser, 1958), i.e. a solution for which factor coefficients are, if possible, either high and thereby indicative of a functional relationship, or else, close to zero. At first glance, the data structure in the current study does not suggest a simple structure besides a functional relationship with the target variable for *all* scores, as all scores measured the same quantity. However, the scores varied in their proportion of noise, and, therefore, varied in the correlation with the target variable. Rotated PCA solutions might have performed that well because they created a simpler data structure by partial neglect of the scores that were least predictive of the target variable. In contrast, the performance of the mean of the six scores suffered from the noisiest scores. Following this interpretation, the potential advantage of rotated PCA solutions, strictly speaking, does not come from a de-noising process but, instead, from a filter process that diminishes the influence of the variables with the highest noise.

### 2.5 Additional posthoc experiments - variations of experiment 2

Experiments 1 and 2 have uncovered some limitations of principal component analysis to infer fundamental cognitive processes from a battery of neuropsychological assessments. Still, across all conditions, the majority of PCA factors after varimax or promax rotation succeeded to identify the fundamental target variables and did so with high precision. In additional posthoc experiments, I varied the experimental conditions to potentially disrupt the performance of rotated PCA factors to identify possible critical limitations of the method.

#### 2.5.1 Experiment 2c – Rare dissociations between target variables

Rotated PCA factors performed outstandingly well in Experiment 2b, even though the target variables were dependent. The statistical dependence of both variables was introduced by the definition of a dissociation pattern. In experiment 2c, I modified the dissociation pattern so that the target variables correlated even higher. Instead of 35 patients with both/no deficit and 15 patients with only one individual deficit each, I changed the groups to a 45-5-5-45 dissociation pattern. With dissociations being that rare, both target variables correlated strongly with r = .63 (SD = .04). The unrotated factors failed almost always – in 98 out of 100 simulations – to align with the target variables, and the Kaiser criterion failed in 48 simulations to find a two-factorial solution. However, rotated factors still performed outstandingly well and consistently succeeded to assess the target variables with high precision. Detailed results are reported in table 3.

#### 2.5.2 Experiment 2d – Removing the ‘anchor’ for rotated solutions

A possible explanation why rotated solutions almost always performed so well could be the presence of a few scores that correlated strongly with each target variable, or at least one score that did so. When factors were rotated, a resulting factor might have been constituted exactly by those scores with high loadings, and therefore assessed the target variable underlying the scores. In other words, one or a few scores out of six could provide an anchor for factor rotation. In experiment 2d, I modified experiment 2b to remove the possible anchor for the second target variable. Again, the first three scores were simulated as a function of the first target variable. The next three scores, however, were simulated as a function of both target variables to the same extent. Therefore, scores that primarily measured the second target variable did not exist.

This simple modification indeed disrupted the performance of PCA to measure the target variables. While 15 out of 100 solutions with unrotated factors failed to align with the target variables, it happened in 57 simulations with the varimax rotation and in 49 with the promax rotation. In the remaining successful rotations, the correlation of the first two factors was numerically lower than before. Additionally, I investigated the performance to measure the first versus the second target variable. For the first target variable, the average correlation with the corresponding factor was still very high (varimax r = .97, promax r = .97). But, as expected, it was much lower for the second target variable (varimax r = .79, promax r = .79). The Kaiser criterion failed to identify the two-factorial structure in all 100 simulations. Detailed results are reported in table 3.

### 2.6 Factor Analysis

Factor analysis was computed for all sub-experiments in experiments 1 and 2. The detailed results are reported in the supplementary. The performance of FA in identifying the underlying variables was not markedly different from PCA. Factor analysis also succeeded to estimate the target variables with good up to almost perfect precision in most conditions. It also occasionally failed to align the factors with the target variables. I decided to not directly compare PCA and FA, as a meaningful comparison would have been limited by the different number of valid factor solutions. On a numerical level, FA was more often able to correctly align the factors with the target variables than PCA, but, on the other hand, the precision was often slightly lower. This was especially the case in the situations with the most challenging data, and, likewise, orthogonal varimax rotation was found to be significantly inferior to oblique promax rotation in these conditions. The average Kaiser-Meyer-Olkin index ranged between 0.67 and 0.88, indicating that the data were suited for factor analysis. Interestingly, this index was lowest in experiment 1b, where the precision of FA to identify the target variables was numerically the best.

### 2.7 Data availability

All the study’s Matlab scripts are publicly available at Mendeley Data (http://dx.doi.org/10.17632/vsg79xx3k8.2). They were used with a pre-defined random number generator seed, hence all data and results can be perfectly replicated.

## 3 Discussion

The present study created a mixed picture of the use of principal component and factor analysis to identify fundamental cognitive processes as it highlights some challenges in the inference of cognitive functions under dependence of data. In principle, PCA and FA seem to be able to infer fundamental cognitive functions with high precision even when variables are highly dependent. However, its success is not guaranteed and depends on certain features of the data.

### 3.1 What can principal component or factor analysis infer from associations in neuropsychology?

For good reasons, neuropsychologists in the past have been suspicious of associations between neurological deficits (see Shallice, 1988). What can we infer from a strong correlation between spatial neglect and hemiparesis of the upper limb of r = .51 (Sperber et al., 2020)? The correlation does not imply that both behavioural measures rely on shared cognitive functions. Both deficits could constitute cognitively and anatomically distinct entities that simply arise from stroke to the same arterial branch. Anyway, a principal component analysis explains 75% of the variance in both variables by a single factor. It would be rash to interpret this factor to represent a fundamental cognitive function. In this context, it seems essential to discuss the two possible meanings of ‘independence’ between cognitive functions. First, functions can be functionally and anatomically independent. This concept is the focus of our research endeavour – we want to identify the distinct, fundamental functions underlying behavioural pathology. Second, two functions can be statistically independent. Statistical dependence, i.e., the association between variables, provides the foundation for PCA or FA to reduce the dimensionality of data. Statistical independence of neuropsychological deficits will rather be the exception than the norm. Brain lesions after stroke follow the vascular architecture of the brain and, thereby, are highly systematic (Zhao et al., 2020). Whenever a lesion induces a cognitive deficit by damage to a specific brain region, the probability of damage to other regions will not be the same anymore. If a lesion damages the left corticospinal tract and causes hemiparesis, it is more likely that the lesion also causes aphasia, as the neural correlates of aphasia are situated in the same arterial territory and close to the corticospinal tract. Moreover, the dependence of deficits is not restricted to neural correlates that are neighbours. Cognitive deficits can be negatively correlated with damage to brain regions that are located further away (Sperber, 2020). For example, if a patient suffers from visual field defects after occipital damage to the primary visual cortex, it is very unlikely that the patient also suffers from frontal lobe damage. Hence, visual field defects and frontal lobe syndrome will negatively correlate. This results in strong associations between most post-stroke deficits (see Figure 3 in Bisogno et al., 2021).

Principal component analysis is mainly a descriptive method (Jolliffe & Cadima, 2016) that can be used to simplify large data or, potentially, gain a deeper understanding of the data. When interpreting the factors that resulted from a PCA, we can try to find a meaningful concept behind them. However, a factor does not necessarily capture an actual entity that exists out there in the world. A striking example comes from the investigation of lifestyle risk factors for cancer, where a PCA factor loaded on smoking and alcohol consumption (Navarro Silvera et al., 2011). This factor can be interpreted in the context of psychological or sociological phenomena. However, the existence of such a factor does not imply the existence of cancer-inducing, nicotine-flavoured beer.

Nonetheless, the present study showed that PCA on neuropsychological data is not restricted to descriptive use, but can infer fundamental cognitive processes. After rotation, the PCA factors were most often able to measure the target variables and, despite the dependence of variables and noise, they often did so with outstanding precision. However, the success of PCA stands and falls with the success of factor rotation, and it is difficult to say if a resulting factor structure failed or succeeded in inferring fundamental cognitive functions. In any case, the intuitive interpretability of factors should be consulted as a plausibility benchmark. In addition, findings from single case studies offer the potential to complement and validate the results of a PCA.

The use of PCA to identify basic cognitive functions closely resembles its use in the mere description of the dimensionality and complexity of clinical profiles. The critical difference between these strategies lies in the interpretation of PCA factors. Whereas the latter assumes factors to represent underlying fundamental cognitive functions, the former is descriptive and assumes factors to represent typical clusters of behavioural deficits (e.g., Azouvi et al., 2002; Zandieh et al., 2012; Corbetta et al., 2015; Sperber & Karnath, 2016). Given its relatively minimalistic assumptions, such interpretation appears to be generally applicable, but it is only of minor relevance to basic cognitive research. A good example for such findings bearing relevance for basic cognitive research comes from the field of spatial neglect, where a multi-factorial PCA solution provided strong evidence that commonly used diagnostics do not measure a unitary syndrome (Azouvi et al., 2002). However, a descriptive interpretation might bear more clinical relevance. For example, the relationship between limb apraxia and aphasia after stroke has been an enigma already for decades, and cognitive scientists have provided many diverging theories if and how both functions share fundamental cognitive processes (e.g. Goldenberg & Randerath, 2015). All this is irrelevant for a neuropsychological therapist. She knows that apraxia and aphasia fall into a typical cluster of post-stroke deficits and often co-occur (Kertesz & Ferro, 1984) – which is merely descriptive information. Therefore, she is aware that therapy should not rely on the compensation of speech production by gestures, which are typically deficient in limb apraxia.

### 3.2 Factor analysis versus principal component analysis

Principal component and factor analysis are mathematically different, however, they are often used in a conceptually similar manner (e.g. Velicer & Jackson, 1990; Field, 2009; Santos et al., 2015). From a mathematical perspective, the interchangeable use of both methods can be questioned, but, still, their results often converge (Velicer & Jackson, 1990). A concept that is often used to differentiate PCA and FA is that the resulting factors constitute manifest factors in the case of PCA and latent factors in the case of FA (Velicer & Jackson, 1990; Velicer & Fava, 1998). The concept of latent factors, which are defined as unobserved, underlying, error-free variables, suits the simulated target variables in the present study. Still, both PCA and FA most often successfully identified the target variables in the present study, and, likewise, both struggled under certain conditions. Both methods sometimes failed to align the first two factors with the target variables and, thereby, critically failed their task. However, such failures were more numerous in PCA. All in all, it appears that both methods are suited to handle the association problem in neuropsychological data with similar limitations. The strong focus of recent neuropsychology on PCA in the identification of latent variables that constitute fundamental cognitive functions (e.g. Halai et al., 2017) does not seem to be an issue. On the other hand, FA was found to be better suited to analyse neuropsychological data from older individuals (Santos et al., 2015). Following this example, real neuropsychological data from patients with brain damage should be used in the future to directly compare the methods with high ecological validity.

### 3.3 The importance of factor rotation

The primary rationale behind factor rotation is to increase the interpretability of factors (Thurstone, 1947; Fabrigar et al., 1999; Field, 2009). The general importance of factor rotation in analyses with psychological data is widely known (e.g. Fabrigar et al., 1999; Sass & Schmitt, 2010; Santos et al., 2015). However, its role may go beyond the mere interpretability of the factors and even affect the construct validity of a factor structure (Sass & Schmitt, 2010), which was mirrored in the present study. In almost all experimental conditions, factor solutions after varimax or promax rotation were superior to the unrotated solution, and the success of PCA or FA to precisely identify the target variables could largely be attributed to this effect. Unrotated factors often even failed to assess the target variables. The success of factor rotation, however, appears to be fragile. In experiment 2d, simple and realistic modifications of the simulations markedly disrupted the performance of rotated solutions. In this situation, the test battery lacked variables that primarily measured the second variable and, hence, no variables existed to anchor the second factor in the rotated solution. In reality, it will be difficult to test if the structure of neuropsychological data allows for a meaningful factor rotation. With a priori hypotheses on possible fundamental functions in mind, one could ensure that the test battery, for each hypothesised function, includes test items that primarily measure this function. Further, a critical and transparent evaluation of the resulting factors for plausibility should be part of any analysis.

A key difference between varimax and promax rotation is that varimax rotation creates independent, orthogonal factors, while promax rotation allows oblique, dependent factors. The orthogonal rotation has often been criticised for being unfit to describe real data, in which variables are often correlated (e.g., Cattell & Dickmann, 1962). In the present study, I found no marked differences between both types of rotations in PCA – even in experiments with correlated target variables the differences between both were non-existent or, at best, negligible. On the other hand, promax rotation was indeed found to be superior to varimax rotation in FA. This was the case in the experiments that were intended to provide the most difficult challenges for the algorithms, i.e. experiments 2c and 2d. This corresponds to previous findings on the superiority of oblique rotation in the description of neurocognitive test data in older individuals (Santos et al., 2015).

### 3.4 The dimensionality of cognitive deficits

Principal component analysis always provides the same number of factors as there are input variables. Only in a second step, a researcher has to decide on a smaller and ideally meaningful set of factors to describe the data. The Kaiser criterion (Kaiser, 1960) is a simple and popular method to estimate the number of meaningful factors. In the current study, the Kaiser criterion often failed to capture the true structure of the data. In all variations of experiments 1 and 2, two target variables existed. However, across experiments, the Kaiser criterion suggested a uni-factorial solution and thereby critically underestimated the dimensionality of cognitive deficits in between 12-100% of all simulations. A first, general reason for this failure might be the limitations of the Kaiser criterion. This criterion is simple to use and often included in statistics packages, but it can lead to seemingly arbitrary decisions and both over-or underestimate the number of factors (Ledesma & Valero-Cabre, 2007; Velicer & Jackson, 1990). Other strategies to estimate the number of factors exist. A similar criterion is the Jolliffe criterion (Jolliffe, 1972), according to which factors with eigenvalues > 0.7 should be retained, which could to some degree counter the Kaiser criterion’s underfactoring found in the present study. Horn’s Parallel Analysis (Horn, 1965) computes factor-wise eigenvalue cutoffs based on a permutation procedure. Could these be used to precisely estimate the number of fundamental cognitive processes underlying neuropsychological test data? Not necessarily. A second, more specific reason possibly exists for the failure of the Kaiser criterion. Cognitive deficits often correlate, and, therefore, even a large number of dissociating cognitive deficits might be describable by a low-dimensional factor solution (see Corbetta et al., 2015; Bisogno et al., 2021). Most criteria to estimate the number of factors, however, aim to find factors that explain considerable variance in the data, i.e. factors with high eigenvalues. In conclusion, a low-dimensional data structure of neuropsychological data does not imply that the underlying cognitive architecture must be low-dimensional. This is already a known issue for normal psychological data, in which unifactor behavioural could result from multiple processes that function in unison (Bejar, 1983). Importantly, in neuropsychological data, this conclusion goes further, as multiple processes that do not operate in unison could be damaged by typical patterns of brain damage in unison.

When we want to infer fundamental cognitive functions, a theory-guided strategy to estimate the number of factors in neuropsychological data could be an alternative. If we already assume that a neuropsychological test battery measures two fundamental functions, we can simply rotate two factors without the consideration of any selection criteria. In the absence of strong a priori hypotheses, we could resort to an exploratory approach. If the addition of a factor does not markedly increase the cumulatively explained variance by the solution, but still yields an intuitively interpretable solution, then this factor should be included. Exploration of possible dissociations in the data, for example for variables that load on the same factor, might provide further guidance.

### 3.5 Principal component analysis to denoise data

Tremendous effort is made to improve the statistical power of lesion-deficit inference methods and the prediction of neuropsychological deficits from brain imaging and clinical data. However, the quality or, more specifically, the reliability of the measured behavioural data set a limit to any use. No matter how sophisticated imaging features and predictive algorithms are, they will inevitably fail to precisely predict a noisy clinical outcome variable. This is a challenge in neuropsychology. Behavioural measures in elderly neurological patients that suffer from deficits across multiple functional domains, including language, motor abilities, attention, and praxis, are often inherently noisy.

Principal component analysis provides a means to improve the reliability of behavioural measures. First, it can create a compound measure from multiple behavioural measures, similar to the arithmetic mean. The current study found no marked differences between a PCA across multiple measures and the mean of these measures in assessing an underlying variable. In this context, any improvements to the reliability of the final measure are to be attributed to the repeated measurement of the same quantity, and not directly to the PCA. However, the computation of the arithmetic mean of multiple variables comes with the tacit assumption that all measures are equivalently scaled. This might be a daring assumption for many neuropsychological instruments. In contrast, PCA does not make assumptions about variable scaling, and might therefore be preferable.

The current study found another benefit of PCA in the measurement of quantities from multiple noisy variables. After factor rotation, the resulting first factor outperformed both the arithmetic mean and the unrotated PCA solution in conditions with high noise. Factor rotation aims to achieve a simple coefficient structure (Kaiser, 1958), where the contribution of the variables to each factor is either high or close to zero. The more noise a variable contains, the lower should be its coefficient for the relevant factor in the unrotated solution. Therefore, a rotated solution might filter out the noisiest items, and, instead, create a compound measure only from those variables that provide a more precise measure of the target variable. It should be noted that this application of PCA is not specific to neuropsychological data in patients with brain injury, which was the focus of the present study but can be applied generally to normal psychological and other data.

The present study simulated random, independent noise that affected variables. In reality, we might be able to trace influences on our variables back to specific, dependent factors. For example, we might realise that our behavioural measures are not just randomly noisy, but systematically affected by working memory capacity. In this example, another fundamental cognitive process comes into play, and the situation resembles experiments 1 and 2. And as discussed above, PCA might again be suited to obtain a precise measure of fundamental cognitive functions. Still, a plausibility test should be performed to validate if we created a more reliable behavioural measure. A comparison with external criteria might be an option. Else, in anatomo-behavioural studies, we can assume that a more reliable behavioural measure should generate higher anatomo-behavioural correlations and models that provide better predictions.

### 3.6 Limitations

The current study’s most important limitation is the use of synthetic data. Real neuropsychological data are more complex than any simulation could be, and, hence, the ecological validity of synthetic data is limited. However, any substantial issues that are found with idealised data in the artificial setting will likely also exist in actual data. On the other hand, synthetic data have the advantage over purely theoretical-methodological work in that they can evaluate methods under specific and controlled conditions. This was especially relevant in the present study, which primarily aimed to evaluate principal component and factor analysis in situations of dependence between variables, which are typical for neuropsychology. Further evaluation of PCA and FA on actual neuropsychological data could provide further insights into this matter.

Another limitation is that principal component and factor analysis with maximum likelihood estimation are not the only methods to reduce the dimensionality of multiple variables. Likewise, alternative strategies to estimate the number of relevant factors or to rotate factors exist (see Sass & Schmitt, 2015). Any method that is based on associations between variables will likely suffer to some degree from the limitations found in the present study. However, it might be possible that other methods for dimensionality reduction and factor rotation are better suited to handle neuropsychological data.

## 4 Conclusions and perspective

Even though principal component and factor analysis are associational methods, they are capable of precisely inferring fundamental cognitive functions from neuropsychological group data. The possibility of false results and other pitfalls, however, puts limitations on this method. In comparison with the vast body of theoretical and empirical methodological works on the dissociation method, I believe that the use of principal component or factor analysis in neuropsychology is still built on an insufficient theoretical foundation. Clear criteria on their use, interpretation, and validation are lacking and, in the face of the increasing number of publications that use the method, this appears to be a pressing matter.

## Supporting information

Supplementary Factor Analyses

