## Supplementary Factor Analyses for "The use of principal component and factor analysis to measure fundamental cognitive processes in neuropsychological data"

Christoph Sperber

### Supplementary

### Additional Factor Analyses

### 1 Methods

With additional analyses, I aimed to evaluate the ability of factor analysis (FA) to handle the association problem in neuropsychological data and infer fundamental cognitive functions. Factor analysis was computed using the ‘factoran’ Matlab function with maximum likelihood estimation. As with principal component analysis (PCA), factors were rotated either with orthogonal varimax or oblique promax rotation. With the predefined random number generator seed, I ensured that the factor analyses were applied to the same simulated behavioural data as principal component analysis. Again, two factors were rotated in all analyses, independent of any criteria to select the number of factors. Additionally, the Kaiser-Meyer-Olkin Index (Kaiser & Rice, 1974; Trujillo-Ortiz et al., 2006) was computed. This measure ranges between 0 and 1 and indicates if a dataset is suitable for a factor analysis. Commonly, values above 0.5 are expected to be required for an adequate factor analysis. The performance of PCA and FA was not directly compared by statistics due to the different numbers of successful solutions. A method that generates more viable solutions (i.e. correctly aligns the two factors with the target variables) might especially do so in simulations that are difficult to handle for the algorithms. Therefore, the overall performance as measured by the correlation between factors and target variables might be worse, thereby preventing a direct comparison of both methods.

### 2 Results

#### *2.1 Simulation experiment 1a – two independent cognitive functions*

The average Kaiser-Meyer-Olkin Index was 0.85 (minimum 0.79), indicating a meritorious degree of common variance for factor analysis. In nine simulations with unrotated factors and eight simulations with promax rotation, the first two factors were not aligned with one target variable each. After the removal of these cases, the factors correlated highly with the target variables. The unrotated solution ($r = .85$) was worse than both rotated solutions, and promax rotation ($r = .93$) was slightly superior to varimax rotation ($r = .96$). Detailed results are reported in supplementary table 1.

#### *2.2 Simulation experiment 1b – two independent cognitive functions and independent test scores*

The average Kaiser-Meyer-Olkin Index was 0.67 (minimum 0.47), indicating a mediocre degree of common variance for factor analysis. The first two factors were misaligned with the target variables in four simulations for unrotated factors and one simulation each for varimax and promax rotation. Promax and varimax rotation (both r = .96) performed almost perfectly and better than the unrotated solution. Detailed results are reported in supplementary table 1.

| Experiment | KMO | Correlation r Factor(s)-Target Variable(s) | | | Significance p (Bonferroni-corrected) | | |
|---|---|---|---|---|---|---|---|
| | | Unrotated | Varimax | Promax | UR-VM | UR-PM | VM-PM |
| 1a | 0.85±0.03 | 0.85±0.08 | 0.96±0.04 | 0.93±0.05 | **< .001** | **< .001** | **< .001** |
| 1b | 0.67±0.07 | 0.93±0.07 | 0.96±0.05 | 0.96±0.05 | **< .001** | **< .001** | = 1.00 |
| 2a | 0.88±0.02 | 0.84±0.10 | 0.95±0.04 | 0.95±0.04 | **< .001** | **< .001** | = 1.00 |
| 2b | 0.71±0.05 | 0.90±0.09 | 0.94±0.06 | 0.96±0.05 | **< .001** | **< .001** | **< .01** |
| 2c | 0.79±0.04 | 0.84±0.12 | 0.89±0.06 | 0.96±0.04 | **< .001** | **< .001** | **< .001** |
| 2d | 0.84±0.03 | 0.84±0.12 | 0.83±0.09 | 0.87±0.09 | = .70 | = .08 | **< .001** |

**Supplementary Table 1 - Detailed results of all experiments with factor analyses**
The first column reports the Kaiser-Meyer-Olkin (KMO) Index. The following three columns report the correlation of the first and second factor with the corresponding target variable. Simulations in which the first two factors failed to align with the target variables are reported in the text and were not considered here. The last three columns show the results of the statistical comparison between the three different FA solutions. Values are mean±standard deviation. Significant results are highlighted in bold. UR = unrotated; VM = varimax; PM = promax.

##### *2.3 Simulation experiment 2a – two dependent cognitive functions*

The average Kaiser-Meyer-Olkin Index was 0.88 (minimum 0.83), indicating a meritorious degree of common variance for factor analysis. The factors failed to describe one target variable each in 56 out of 100 simulations with the unrotated solution and three simulations with promax rotation. Again, rotated solutions outperformed unrotated ones (both varimax and promax r = .95). Detailed results are reported in supplementary table 1.

##### *2.4 Simulation experiment 2b – two dependent cognitive functions and independent test scores*

The average Kaiser-Meyer-Olkin Index was 0.71 (minimum 0.51), indicating a middling degree of common variance for factor analysis. The first two factors always aligned with one target variable each for rotated solutions. The unrotated solution, however, failed in 18 out of 100 cases. Both rotated solutions outperformed the unrotated solution, with promax rotation (r = .96) being slightly better than varimax rotation (r = .94). Detailed results are reported in supplementary table 1.

#### *2.5 Experiment 2c – Rare dissociations between target variables*

The average Kaiser-Meyer-Olkin Index was 0.79 (minimum 0.62), indicating a middling degree of common variance for factor analysis. The unrotated factors failed in 70 out of 100 simulations to align with the target variables. Rotated factors still performed well and succeeded to assess the target variables with high precision in 98 out of 100 simulations. Notably, promax rotation (r = 96) outperformed varimax rotation (r = .89). Detailed results are reported in supplementary table 1.

#### *2.6 Experiment 2d – Removing the 'anchor' for rotated solutions*

The average Kaiser-Meyer-Olkin Index was 0.84 (minimum 0.73), indicating a meritorious degree of common variance for factor analysis. Unrotated solutions failed to align with the target variables in 37 out of 100 simulations, after varimax rotation in seven simulations and with promax rotation in 37 simulations. As for PCA, the correlation of the first two factors after rotation was numerically lower than before. Additionally, I investigated the performance to measure the first versus the second target variable. For the first target variable, the average correlation with the corresponding factor was still decent up to almost perfect (varimax r = .88, promax r = .96) with an advantage of promax rotation. But, similar to PCA, the precision was lower for the second target variable (varimax r = .78, promax r = .79). Detailed results are reported in supplementary table 1.

#### *Additional References*

Kaiser, H. F., & Rice, J. (1974). Little Jiffy, Mark Iv. *Educational and Psychological Measurement, 34*(1), 111–117. https://doi.org/10.1177/001316447403400115

Trujillo-Ortiz, A., R. Hernandez-Walls, A. Castro-Perez, K. Barba-Rojo and A. Otero-Limon (2006). kmo:Kaiser-Meyer-Olkin Measure of Sampling Adequacy. A MATLAB file. [WWW document]. URL http://www.mathworks.com/matlabcentral/fileexchange/loadFile.do?objectId=12736
